# Information theoretic inference of magnitude and direction of gene flow in metapopulation networks using nyemtaay, with potential for applications in metastasizing cancer clonal cell origin analysis

**DOI:** 10.1101/2024.06.04.596026

**Authors:** Adrian N. Ortiz-Velez, Jeet Sukumaran

## Abstract

**Background:** We introduce `nyemtaay`, a Python package for the calculation of classical population genetic statistics and inference of gene flow network connections and directionality in metapopulation networks using information theory. This genetic information flow network inference approach provided here is the only existing implementation of [1], and is applicable not only to ecological and evolutionary organism and landscape scale studies, but also has potential applications in, for example, cancer biology for analyzing clonal cell origins in metastasizing tumors.

**Results:** We demonstrate this potential through simulations and an analysis of metastasizing cancer cell lineages, showcasing its ability to identify the tissue site of origin in cancer networks. This work highlights the importance of considering demographic history and founder effects in interpreting gene flow directionality, and the benefits of this understanding in allowing application of this approach to gene flow network modeling to reach a broader range of domains.

**Conclusions:** `nyemtaay` is available under the MIT license from its public repository (https://github.com/aortizsax/nyemtaay), and can be installed locally using the Python package manager ‘pip’.

## 1 Introduction

Here, we present `nyemtaay`^1^, a Python package that in addition to providing a suite of ## traditional population genetic metrics such as nucleotide diversity, *F*_*st*_, etc., also provides a novel implementation of information theoretic divergence statistics [2, 3] that support inference of gene flow direction in networks of genetically-related populations [1]. As with all information theoretic approaches, the metric extracts signal from patterns in the data without reference to any mechanistic or phenomenological processes, models, statistics or assumptions. This provides potential broad applicability to many different systems, with previous applications to networks of populations related through concurrent gene exchange at classical population genetic landscape scales [4] as well as in other information spaces with similar architectures, biomedical applications at microbiotic scales (cancer clonal systems), as well as to populations distributed in time and/or space, such as those related by historical gene exchange relationships (e.g., Neanderthals and humans). While pattern-based approaches do not have any particular model or generative process assumptions, however, the expressions that we use to extract information from data, and the interpretation of various measures based on the information, may require evaluation and possibly modification in applications where other processes may generate patterns that also structure the data. In the example here, we show that, while the statistic is effective in diagnosing asymmetry in general in the context in which it was developed, correctly interpreting the specific directionality requires taking into consideration of the demographic history of the populations in addition to their relationships, specifically with respect to processes or mechanisms by which genetic diversity can be reduced such as bottlenecks, founder effects or genetic drift. We demonstrate this using simulations, and demonstrate usage of the site frequency spectrum as a heuristic tool to identify a reduction of genetic diversity in founder effect populations. We then provide a sample analysis of a published known network of metastasizing cancer cell lineages to identify the tissue site from which the network originated and compare the results.

## 2 Materials and Methods

### 2.1 Package description

`nyemtaay` is a pure Python package for the analysis of relationships of genetic samples in a population genetic framework that features the ability infer the gene flow network connectivity between sampling demes and any directionality in the connections.

### 2.2 Classical population genetics analyses

The classical population genetic statistics provided by `nyemtaay` are described in detail in the appendix. While these statistics are available in a variety of other packages in other languages, `nyemtaay` provides independent implementation which are useful for testing/referencing purposes, as well as simplifying dependency stacks.

### 2.3 Information theoretic directed gene flow network inference

`nyemtaay` organizes information extracted from a collection of genetic samples into a *directed genetic information flow network*, using an approach described by [1, 4]^2^, in two phases: by first (a) inferring the network structure in terms of pairwise node connections and the relative magnitude of the information flow between them based on entropic measures of genetic differences; followed by, (b) inferring a net directionality for each pair of connected nodes based on calculating and then interpreting an entropic index of segregation.

#### 2.3.1 Information flow network inference

Following [1, 4], given a collection of samples that can be organized into nominal clusters based on some *a priori* understanding of affinities (for e.g., the samples are associated with particular geographical or tissue sites), `nyemtaay` implements the calculation of the normalized Jensen-Shannon divergence (JSD) [5] between each distinct pair of clusters based on measures of information [2] extracted from genetic sequences in the samples from each of the clusters, and then constructs a gene flow network by connecting nodes representing the clusters based on the percolation threshold of the network [1, 6].

The entropy and mutual information [3] of the clusters are extracted from the genetic sequences based on patterns of relative frequencies of shared vs. exclusive alleles in the samples from both clusters [1, 4]. These measures of information facilitate the calculation of relative entropy, also known as the Kullback-Leibler (KL) divergence [7], between any two clusters. The Jensen-Shannon Divergence (JSD) between any pair of clusters is then given by the weighted average of the KL divergences of each cluster from their mutual mean 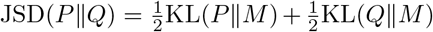, where 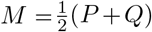 is a mixture distribution that is the average of the distributions *P* and *Q* (in this context, the pooled allele frequencies). The JSD is an entropy-based [2] measure of difference between two probability distributions, as is the KL divergence on which it is based. Unlike the KL divergence, however, the JSD is bounded between 0 and 1, and is symmetric (though still not a metric or true distance due to not fulfilling the triangle inequality axiom; the square root of this, however, does fulfill this and is a true distance). The JSD can be use to compare probability distributions with non matching supports [4, 8]. The JSD can be interpreted as providing a measure of divergence between the allele frequencies of each of the clusters being compared, taking into account patterns of alleles that are shared across clusters as well as those that are unique to one or the other cluster. These JSD values are then normalized to account for varying population size [1]. The normalized JSD values between each distinct pair of clusters are used to construct a gene flow network between the clusters. Each pair of clusters are connected to each other with the a direct gene flow path if their a mutual JSD value is equal to or below the percolation threshold value for the system. The percolation threshold describes the point at which a system transitions from a state where there are small isolated clusters to one where there exists at least one spanning cluster, i.e., a connected component that provides a continuous path across the system [9]. In this application, as we increase the divergence value below above which we assume clusters are isolated, the percolation threshold is the minimum divergence value at which a path emerges through the network that spans the entire area [1]. All distinct pairs of clusters that have JSD values below this threshold are considered to have sufficient convergence in genetic information to indicate gene flow, while those above the threshold are considered to be sufficiently divergent in genetic information to indicate lack of direct gene flow.

This approach to gene flow network inference — with direct connections between clusters indicated by entropy-based divergence values equal to or below the percolation threshold, with divergence values on the basis of differences in information measurements of allele frequency patterns — has been shown to effectively replicate population and metapopulation structure [1, 6].

#### 2.3.2 Information flow direction inference and interpretation

The JSD value between the two clusters, normalized or otherwise, gives a measure of the magnitude of (genetic) information flow between any two sampling clusters, but as a symmetric value, not any indication of direction. [1] proposed a novel index of information flow between two populations that calculates and takes the difference in the entropies of the allele frequency distributions of each population, but measured and normalized over the joint dominium of their individual distributions. This allows the populations to be ranked in terms of the entropies of their distribution of their allele frequencies, but adjusted so as to only consider shared alleles. The index *I*(*P* → *Q*) = +1 indicates that the information carried by population *P* in the joint allele distribution dominium of *P* is greater than that of the joint dominium of *Q*, and vice versa, if *I*(*P* → *Q*) = − 1. These values are taken to be indicative of gene flow from *P* to *Q* and vice versa on the reasoning that information flows from the distribution which has the lower entropy:”The idea is that the net flow of gametes is from the less entropic (i.e., less diverse, more segregated) populations towards the more entropic, properly adjusting for the different sizes and fraction of common alleles” [1]. The significance of the information flow indication is then tested by randomization approaches [1].

### 2.4 Application to colonization and founder effect demographic history

We are excited about the potential for application of the approaches to gene flow network inference provided by `nyemtaay` not only within the ecological genetics context in which it was developed, but, more broadly, in any context where information flow between populations structures the information within them. Indeed, it has been successfully applied in other domains such as [10, 11], demonstrating the advantages of the model-free and pattern-based approaches that characterizes information theoretic methods in general. However, nonetheless, it is important to ensure that some of the information extraction principles (e.g., the expectation of a symbol against which the information gained by observing it is measured) and interpretive reasoning (e.g. where source populations are identified on the basis of having a lower entropy), are still valid when applied to these other contexts.

#### Demographic history: founder effect information flow direction inference inversion

While successfully providing insight regarding gene flow direction in some very useful applications [1, 6], as noted by [1], there are contexts in which the assumptions underlying the intepretation of result (rather than the assumptions of the calculations) may be violated, leading to incorrect reasoning regarding direction of the gene flow. For example, if an external event or process reduces the genetic diversity in one population, the population will be diagnosed the source population of assymetric gene flow, even when it is in fact the sink [1]. There are a number of processes or mechanisms by which a sink population might present reduced genetic information and thereby require inverted interpretation of the relationship of lower vs higher joint-domina normalized relative entropies of each pair of populations under consideration: bottleneck or founder effects, or more generally any reduction in population sizes such that genetic drift might dominate. One area concern here would be founder effects which would characterize population in systems with recent colonizations or invasion events, as these are very often systems in which gene flow directionality might be important evolutionary biology, conservation, or medical concern — to identify invasive population origins for management or further study, for example — and yet are precisely the systems prone to misinterpretation due to the lower diversity of the derived (founded) populations.

#### Evolutionary vs. ecological gene flow patterns

The *current* theoretical framework for this approach to gene flow network inference was developed in the context of spatial ecological genetics, where all populations are exchanging genes in continuous ongoing gene flow, with no consideration of the evolutionary history of the populations. There has not been any evaluation of the performance of this approach in the latter context, which would characterize a number of systems for potential application.

#### 2.4.1 Baseline case: “historyless” fully-contemporaneous purely-spatial “spanning” gene flow

The original context in which the gene flow network in which the information theory measures were formulated — i.e., how a particular distribution of alleles maps to a quantification of the system’s entropy and the background expected value against which this is measured — as well the interpretation of direction of gene flow based on the differences of entropy was that of a set of distinct contemporaneous populations continuously exchanging genes through migration with no evolutionary or demographic history – a “spanning” rather than “branching” gene flow pattern. The *only* relationships connecting the populations in these contexts are these current, ongoing, and contemporary ones, gene flow through continuous ongoing migration, in contrast to, for e.g., models that include demographic events such as population fragmentation, isolation, or splitting. In contrast to the experimental treatments considered in the next section, the populations have no historical relationship to each other in the sense that one was the ancestral or origin population of the other: a “source” population here is exclusively in terms of current ongoing migration in space, rather than an ancestor in time.

To demonstrate efficacy and assess baseline performance in this case, we simulated data sampled from three populations, *A, B*, and *C*, under two scenarios: (1) a (spatial) “spanning radiation” and a (2) (spatial) “spanning stepping-stone”.

Population sizes in all cases were fixed at *N*_*A*_ = *N*_*B*_ = *B*_*C*_ = 100000 and the mutation rate at 1*e* − 8, while migration rates systematically varied over 1*e* − 4, 1*e* − 5 and 1*e* − 6 as experimental treatments. The `DEMES` [12] package in conjunction with `msprime` [13] was used to generate the data, driven by custom scripts available as part of the `nyemtaay` distribution.

The resulting data was analyzed in `nyemtaay`, with following expected results:

- In the spatial radiation cases, we expect the inferred gene flow network to have paths *A* → *B* and *A* → *C* and not have any other path, in particular not have path *B* → *C*.
- In the stepping stone cases, we expect the inferred gene flow network to have paths *A* → *B* and *B* → *C* and not have any other path, in particular not have path *A* → *C*.

#### 2.4.2 Demonstration of performance under confounding processes

To evaluate the applicability of this approach of estimating gene flow relationships along a temporal axis (ancestor-descendent), and demonstrate the impact of founder effects of on the analysis, we present a simulation study as shown in Figure 2.

**Fig. 1.**
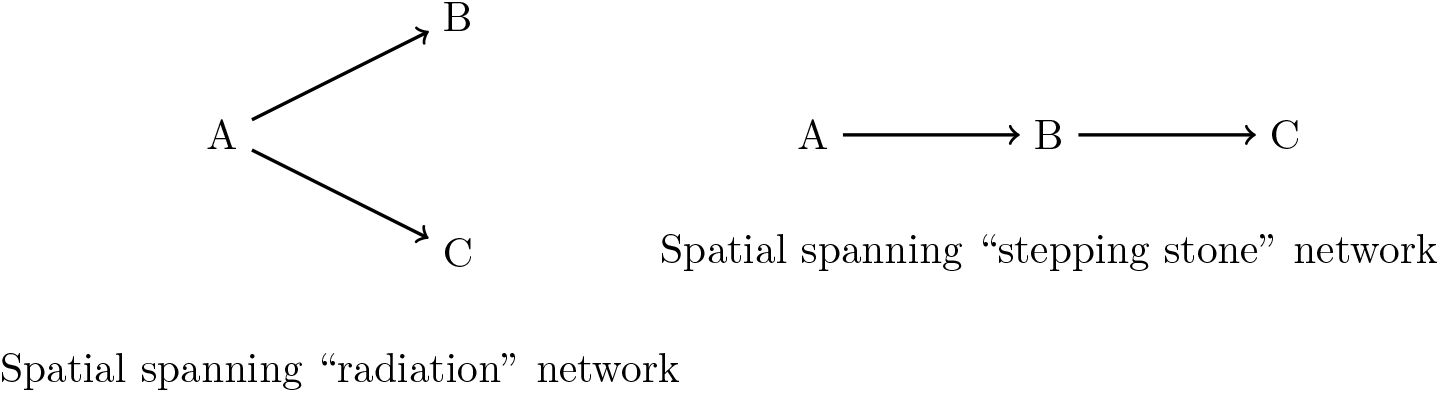
Gene flow scenarios for populations A, B, and C. Left: Spanning radiation scenario with direct flows from A to both B and C. Right: Spanning stepping stone scenario with sequential flow from A to B, then B to C.

**Fig. 2.**
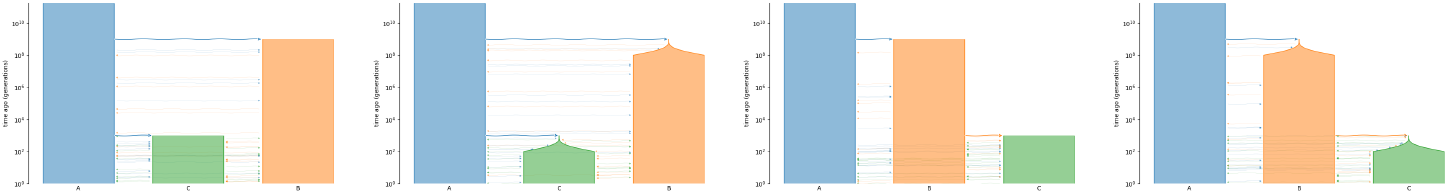
Experimental design to demonstrate impact of demographic founder effects on gene flow network directionality inference using the approach of [1]: (a) the radiation model without founder structuring effects; (b) the radiation model with founder structuring effects; (c) the stepping stone model without founder structuring effects; and (d) the stepping stone model with founder structuring effects. These diagrams are provide the general experimental scenarios, but parameter details (divergence times, founder population sizes, migration rates, etc.) varied across experiment treatment. See text for details.

A *full* assessment of this approach will require far more comprehensive and rigorous experiment design. We could in principle locate regions of parameter space or an experiment design in which this approach succeeds rather than fails, but that will not help any researcher with the particular case than the demonstrative analysis we present here, and moreover, may detract from the message that researcher are obliged to test the approach with simulated data calibrated on their own system before proceeding (a protocol that really should be adopted for *any* method). Our purpose here is to demonstrate to researchers, that, while we advocate application of this approach to other useful problems, until future work develops modifications to the theory and statistics, this broader application calls for case-by-case pre-analytical assessments of suitability using simulation-based performance test calibrated to the systems of interest. The current directionality indicator statistic is one of the simplest possible out of a range of segregation indices, and numerous potential avenues for other quantifiers or indiciators for this remain to be explored, and this should be the focus of future work rather than exhaustively stress-testing a directionality indicator that works well enough when given the context and assumptions under which it was developed. Future work should be directed toward developing a suite of directionality indicators with known behaviorial characteristics that can either be insensitive to unknown evolutionary or demographic effects, or otherwise explicitly take into account these effects.

The basic experimental design involves three populations related through: (a) a “branching radiation” model of evolution, with ancestral population “A” giving rise to population “B” and later on “C”, with all populations surviving to and sampled in the present day; and (b) a “branching stepping-stone” model of evolution, with an ancestral population “A” giving rising to “B”, and “B” later on to “C”, with all populations surviving to and sampled in the present day.

In the “branching radiation” branching model, a sequential series of population “budding” or spawning events connects both “B” and “C” populations as descendent as well as gene flow sinks of population “A”: at time *t* = *T*1 generations in the past population *B* originates as a colony of population *A*, followed by, at time *t* = *T*2 generations in the past, population *C* originates as a colony of population *A*. (2 The “branching stepping-stone” model is similar, except that population “C” arises as a a descendent and persists as a gene flow sink of population “B” rather than “A”: at time *t* = *T*1 generations in the past population *B* originates as a colony of population *A*, followed by, at time *t* = *T*2 generations in the past, population *C* originates as a colony of population *B*. (2. The effective population size of population “A” remains stable through its history at *N*_*e*_. The population sizes of *B* and *C* experience a founder effect at colonization of *N*_*f*_, with growth such that the original population size of *N*_*e*_ is reached after 0.10 proportion of the time since the split has elapsed. So, for example, if a population split from the ancestor *T*1 generations ago with new colony population size of *N*_*f*_, it wil grown to reach the ancestral stable size of *N*_*e*_ by 0.9*T*1 generations ago.

Three different dimensions of impact on performance were explored. First, the impact of age or time since the splitting events was explored by five distinct experimental treatments, each corresponding to different parameters of *T*_1_ ∈ {1*E*4, 1*E*6, 1*E*8, 1*E*10, 1*E*12}, but all with 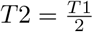. Second, the impact of the founder effect by varying the size of the initial colony populations *N*_*f*_ ∈ {1, 100, 1000, 100000}, corresponding to {1*E* − 5, 1*E* − 3, 1*E* − 2, 1.0} proportion of the ancestral and current population size, with the final treatment here the null case of no founder effect. And, finally, the rate of migration, with three migration rate treatments of 0.0 (total isolation following population splitting), 1*e* − 4, and 1*e* − 6. Migration was included in the systems only to reinforce the relationship signal (i.e., from “A” to “B” and “C” in the radiation case and from “A” to “B” and “A” to “C” in the stepping stone case) to see if this dynamic, as part of the original context in which this approach was developed will improve performance, with a migration rate of 0.0 as a control (i.e., the only gene flow is the from the migrants of the initial colonization pulse).

A total of 30 replicates were run under each experimental design (i.e., all distinct combination of treatments and parameters), and each dataset was analyzed separately using `nyemtaay`.

Expected results are shown in the table below:

**Table 1.** Comparison of expected proportion of counts in each category of directionality indicator values: positive (indicating flow from *i* to *j*, e.g. *A* to *B* if *I*_*AB*_), negative (indicating flow from *j* to *i*, e.g. *B* to *A* if *I*_*AB*_), or zero (path not inferred).

| Path | Splitting Model | Exp $I_{ij} > 0$ | Exp $I_{ij} < 0$ | Exp $I_{ij} = 0$ |
| --- | --- | --- | --- | --- |
| I_AB | Branching-radiation | 1.0 | 0.0 | 0.0 |
| I_AC | Branching-radiation | 1.0 | 0.0 | 0.0 |
| I_BC | Branching-radiation | 0.0 | 0.0 | 1.0 |
| I_AB | Branching-stepping-stone | 1.0 | 0.0 | 0.0 |
| I_AC | Branching-stepping-stone | 0.0 | 0.0 | 1.0 |
| I_BC | Branching-stepping-stone | 1.0 | 0.0 | 0.0 |

#### 2.5 Metastatizing cancer information network flow analysis

Here a demonstrate the *potential* for novel application of this informational theoretic spatial ecological gene flow network inference in the biomedical domain to tracing the origin tissue site of metasizing subclonal cell network. While gene flow analysis has traditionally been considered in the context of collections of reproductive organisms, we can apply the `nyemtaay` approaches at this suborganismal scale of metastasizing clonal and subclonal cell relationships by modeling a metastasizing cell line gene flow network that connects tissues sampling sites to each other, with each tissue sampling site corresponding to a distinct population of genetic information, and genetic information exchanged along the network connections through metasizing cell migration between tissue sites. We can view the trajectory of cancer cell metastasis and spread as establishing a gene flow network connecting the various cancer sites (Quinn et al. 2021, El-Kebir et al. 2018, Lote et al. 2017, Zhao et al. 2016). Each tumor in a metastasizing cancer system, as a colony or population of cells, can be represented as a distinct node in a genetic information flow network network, connected to other such nodes by asymmetrical information flow in time representing the colonization history connecting primary, secondary, tertiary, etc. tumor sites. As the primary tumor initially seeds secondary tumors at other sites in the host through direct movement of cells from the site of origin to other locations in the body, and these in turn each potentially seed their own tumors, an information flow network mapping these dynamics would reflect the primary tumor population as a source and its immediate secondaries as sinks, and so on for the relationship between the secondaries as the source of information for *their* secondary sinks. Following the initial seeding, there might of course be multiple other information flow paths connecting any two of the tumors, including not just bidirectional flow between descendents, but also more complex “backflow” cycles. Theses metastatic and post-metastasis cell movement historoies create the information flow paths connecting the tumor nodes in the cancer metastatic migration network graph. An advantage of using gene flow networks as opposed to phylogenetic approaches to analyzing these relationships is the relatively simplicity of accommodating post metastatic colonization gene flow, while an advantage of using information theoretic approaches is the computational efficiency as well as lack of model or process assumptions.

Here we apply `nyemtaay` to the finer-scale primary tumor data from (Yang et al. 2022). In this study, cell lineage histories within a genetically engineered mouse model are tracked through inheritable indel patterns introduced by the CRISPR-Cas9 system. These indels act as unique barcodes that are transcribed, enabling their detection via single-cell RNA sequencing (scRNA-seq). Phylogenetic approaches were used to construct detailed evolutionary trees of cancer cell populations within tumors, based on inheritable indel patterns introduced via the CRISPR-Cas9 system and captured through single-cell RNA sequencing (scRNA-seq).

Our objective is to demonstrate as proof-of-concept, the applicability of `nyemtaay` to reconstruct the information flow network that connects six tumor sites from which the data were sample: the primary tumor (*T*1), a soft tissue metastatic site (*ST*), and three liver metastatic sites (*L*1, *L*2, and *L*3). Recognizing *a priori* that the demo-graphic dynamics of the metasizing cell lineage networks not only typically include multiple founder effects or bottlenecks, but with rapid proliferation of single cell lineages present extreme founder or bottleneck effects on the genetic information, we predict that the polarity of the interpretation of gene flow direction to be inverted, as discussed above. We note that, a general understanding of the system dynamics is sufficient basis for such a prediction ina proof-of-concept work such as this, in practical applications to new datasets, secondary analyses should applied to determine this, as suggested by [1, 4], and, in cases where some of the information flow processes may be more complex, new theory might need to be developed as well.

## 3 Results

### 3.1 Information flow analysis of baseline spatial population gene flow with no demographic structuring

The results are summarized in the form of an ANOVA analysis of splitting model and migration as treatment factors and visualized in Figure 4. Inference performance as measured by RMSE was optimum at the lowest migration rate, 1*e* − 06, with RSME of less than 1.0, reflecting accuracy rates, as given by proportion of replicates in which a path was recovered with the correct direction, ranging from 0.8 to 1.0. Conversely, performance was worst at the highest migration rate, 1*e* − 04, with RSME rates in the 0.2 to 0.4 range, and accuracy rates below 0.5.

**Fig. 3.**
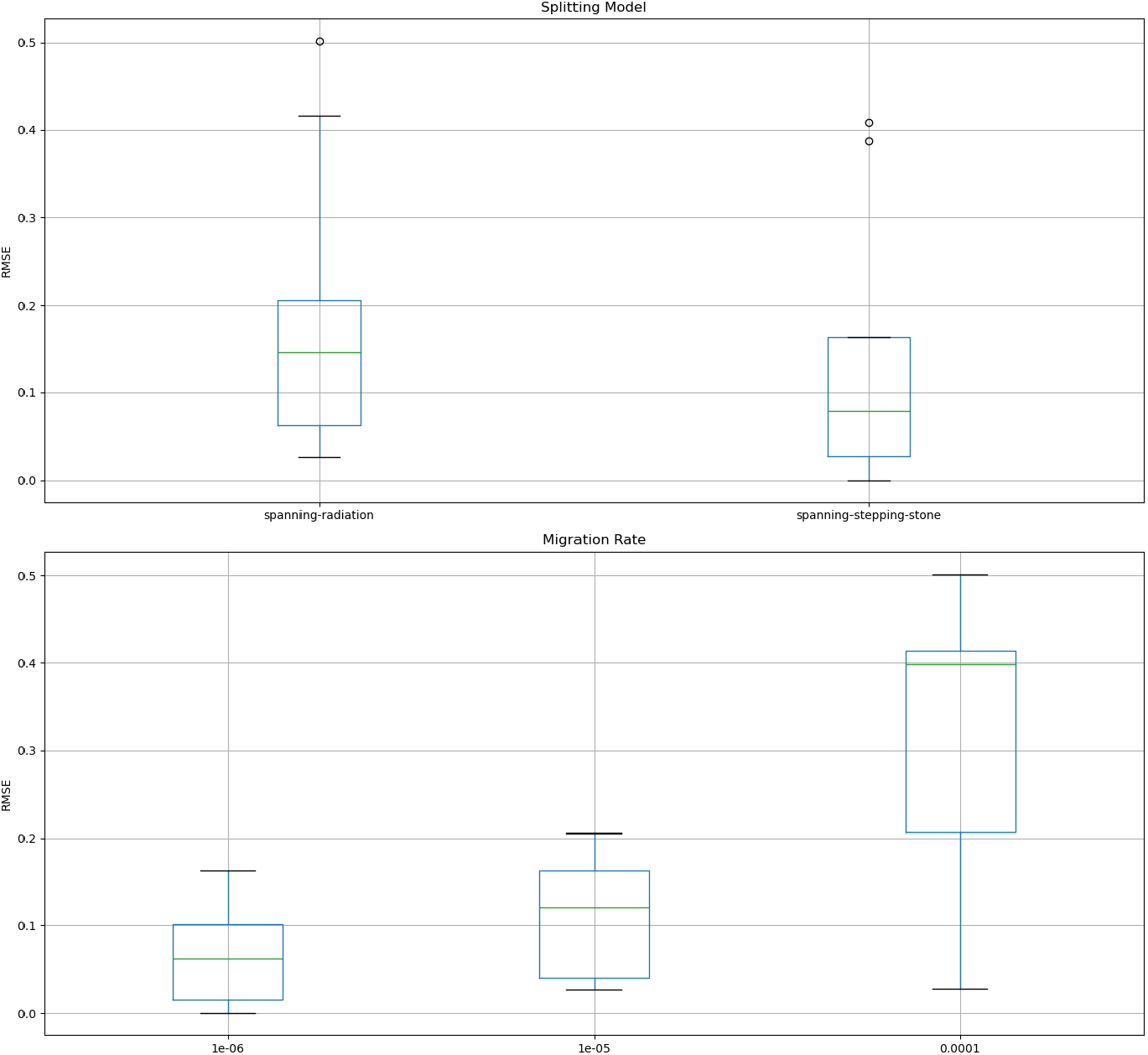

**Fig. 4.**
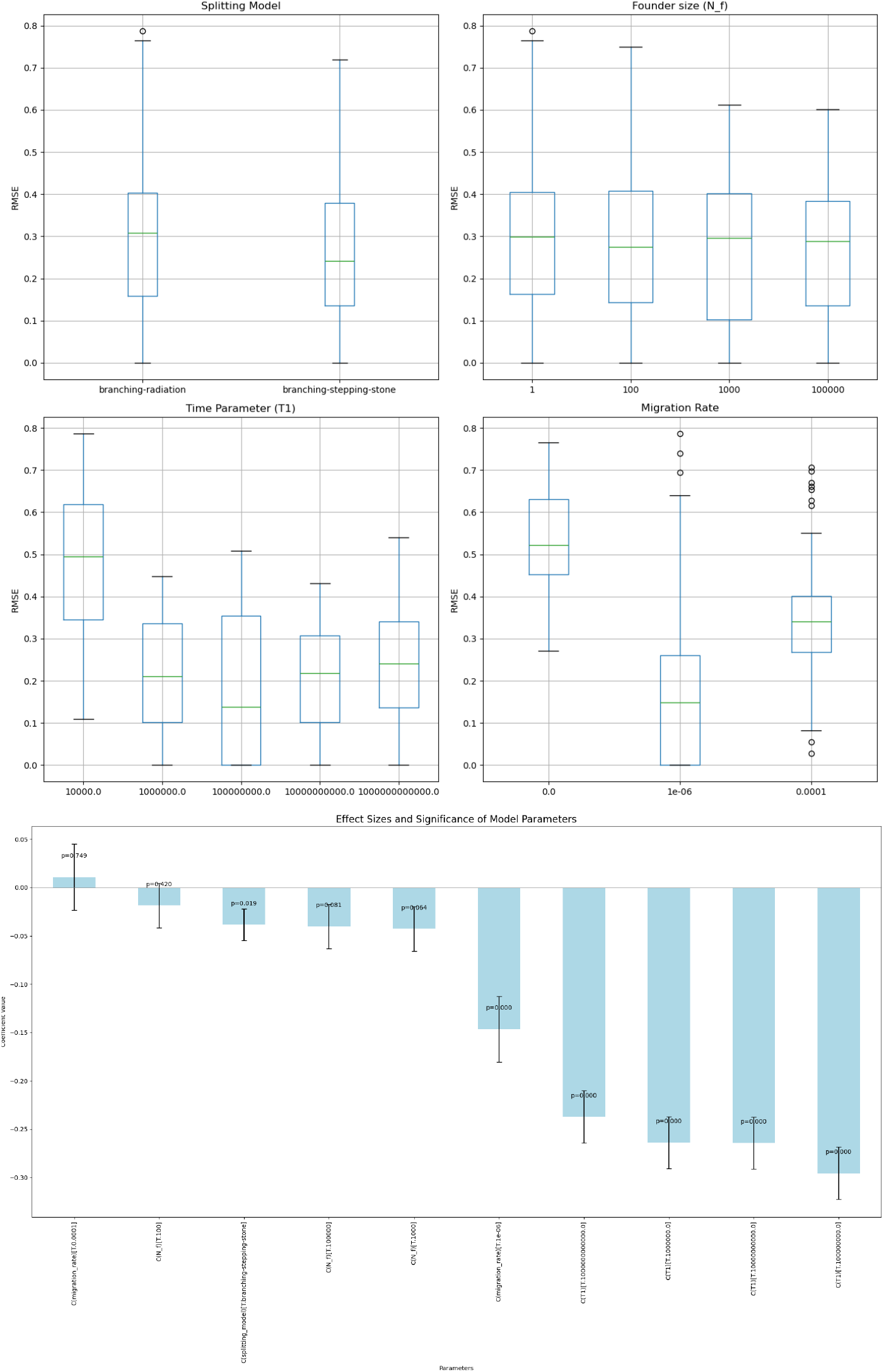

### 3.2 Information flow analysis of serial ancestor-descendent population evolution with founder effects

Here again, the results are summarized in the form of a visualization of the linear model of treatment effects on RSME, showing key parameters of interest (splitting model, *T*1, *N*_*f*_, and migration rate) in terms of their correlation to the RMSE (measured against the predicted indicator values of *I*_*AB*_, *I*_*AC*_, and *I*_*BC*_) as well as an ANOVA analysis in terms of parameter impact is shown in Figure 4. The performance of the inference with gene flow networks structured by evolutionary branching relationships was generally worse than the spanning configurations, except for one particular region of the parameter space in which performance was generally better. The splitting model marginal RSME’s (i.e., the RSME’s of the splitting models summarized over all other treatments), were degraded from the 0.05−− 0.2 range in the spanning cases (Figure 4, top) to the 0.4 −−0.15 range in the branching cases (Figure 4, top left). Founder effects did not have any significant effect (Figure 4, top right), but spanned an RSME range of 0.1−−0.4 regardless of the founding population value. Both the divergence time age and the migration rate showed a “sweet spot” of performance. While shallower split times (1*E*5 generations) had significantly degraded performance (RSME in the 0.35−−0.6 range, deeper time values (≥ 100000 generations) had significantly improved performance as measured by lowered RSME scores (Figure 4, middle left and bottom), with some treatments including performance that even exceeded the spanning cases (e.g., at the moderate split time value of 100000.0). Migration rates also had a major impact on performance, with peak performance in the 1*e* − 6 range and significantly and strongly degraded performance at lower or zero migration rates (Figure 4, middle right and bottom).

### 3.3 Information flow analysis of metastasizing cancer clonal cell network

The original analysis of mouse model cancer system reported two large metastatic expansion events from the primary tumor site *T*1, with the first to the soft tissue site *ST*, and the second to the three liver site *L*1, *L*2, and *L*3. The information flow network inferred by `nyemtaay` recovered this fundamental structure, with one difference: while the primary tumor was linked to both subclonal expansionsand two subclonal expansions had no edges connecting them, instead of a direct connection between the primary tumor and liver site *L*3, `nyemtaay` inferred a secondary spread from *L*2 to *L*3. As noted in the original paper, and as discussed and demonstrated here, bottleneck or founder effects resulting in lowered diversity in the derived sink populations will invert the entropy based index relationships, and, indeed, the directionality indicator recovered exactly the opposite direction in every single edge for which previous edge information was available.

## 4 Discussion

### 4.1 Information flow in spanning vs. branching networks

We showed that, in one particular region of parameter space, across the two basic contrasting 3-node network stuctures examined, the performance of information flow network inference approach provided by `nyemtaay` was degraded when applied to networks with ancestor-descendent relationships between nodes. We have no doubt that we might be able to locate some other equally plausible experimental design in which the performance may be improved or further degraded, but rather than focusing on the performance envelope of the current approach, we adopt the view that a more productive course would be to instead focus future work on identifying the aspects or components of the current approach that need to be adapted. Similarly, while we found that there were regions in which performance was extremely good, on par with the spanning case performance, we emphasize that this should not be taken to mean that the this approach has optimal performance with migrations rates of, for e.g., 1*e* − 6. This characterization is representative of not just a particular part of parameter space considered, including the fixed mutation rates and population sizes, but also the overall experimental design in terms of branching patterns and migration rate treatments.

We report our work here less as broad assessment of this approach across a range of scenarios and conditions in which may be applied, but rather to communicate the understanding that this approach to gene flow network inference, like *any* method, requires caution when applied in contexts in which some of its interpretative structures might be impacted adversely by confounding processes. We also show how using simple simulations we can gain an understanding of which parts of parameter space we may expect optimum performance. We suggest that researchers interested in applying any method, novel or otherwise, for the first time with their own datasets, should consider carrying out simulation-based performance assessment of the method in a region of parameter space centered on their data.

### 4.2 Ecological and evolutionary biological analysis of cancer: relative new frontiers in medicine

`nyemtaay` successfully recovered the information flow connections in a metastatic sub-clonal expansion network of cancer cells, and, with *a priori* understanding of the founder effect characterizing the dynamics of this system, we were able to correctly infer the directionality of the gene flow. We recognize that this understanding may not apply with a range of other systems, whether in macro-, meso-, or microecological domains, and, furthermore, may not even be so simply characterized as either/or — in some systems we might expect some population events to be structured one way while others another. Our contribution toward this direction is, we hope, just part of a growing trend of cross-application of tools developed to study biological processes at one scale being applied to another. There remains much work to be done to support these transitions, and less to do with adapting to the different scale as much as to the greater prominence of confounding signals in the patterns. While in some systems tests for founder effects or bottleneck effects are available, these are only some of the processes that might be involved, and some of the model assumptions might not be applicable or desirable in the contexts, e.g. in metastatic cell network evolution. In this particular case, the simple directionality indexes is based on comparing allele frequencies, and when the genetic diversity of a population is dramatically restructured by drift, as occurs during bottleneck dynamics or founder effects, this reverses the apparent information flow in the system. This is particularly problematical in a number of application domains in which direction of information flow is of special interest, such as invasive or colonization ecological biology, infectious disease dynamics, or, as demonstrated here, in identifying metastatic information flow paths connecting tumors to identify the tumor or tissue sites in which the origin cell is located. More work is needed to: (a) improving the directionality index to account for potential drift effects, alongside (b) understand applicability of population genetic founder effect/bottleneck detection approaches to microecological spaces and systems such as cancer, and adapting them as needed. The benefits of being able to apply these approaches in domains such as these are clear, and development of further theory and statistics to facilitate these remains in the future.

### 4.3 Comparable approaches

`nyemtaay` offers a pattern-based approach to estimation of an important parameter in a number of studies that complements existing methods, sophisticated and powerful as they are. BAYESASS [14], for example, offers many advantages over previous approaches gene flow network inference and analysis, being not constrained by the unrealistically restrictive assumptions of most classical population genetic statistics; in particular, in this case, the Hardy-Weinberg equilibrium. However, due traditional challenges of computational Bayesian inference such as MCMC convergence and mixing [15] and sensitivity to model misspecification [16], as well as the computational expense of a full-likelihood Bayesian approach, there will no doubt be cases where an information theoretic pattern-based such as that provided by `nyemtaay` might be useful for comparison given its advantages and demonstrated success in ecological as well as evolutionary genetics [17**?** –19].

## 5 Availability and requirements

**Project name:** `nyemtaay`

**Project home page:** https://github.com/aortizsax/nyemtaay

**Operating system(s):** Platform independent

**Programming language:** Python

**Other requirements:** Python 3.10 or higher

**License:** New BSD License

**Any restrictions to use by non-academics:** None

## 6 Declarations

### 6.3 Ethics approval and consent to participate

Not applicable.

### 6.4 Consent for publication

Not applicable.

### 6.5 Data availability

The results and analysis reported in the current study are available for download from the code repository: https://github.com/aortizsax/nyemtaay.

### 6.6 Materials availability

Not applicable.

### 6.7 Code availability

The source code, documentation, example data sets, etc. are available for download, installation, or direct usage from the `nyemtaay` code repository: https://github.com/aortizsax/nyemtaay.

### 6.8 Author contribution

AOV: developed the `nyemtaay` code, including native Python implementations of all statistics; identified, downloaded, curated, and analyzed all the empirical datasets; contributed to the writing, ideas, and discussion. JS: conceived of, planned, and mentored the project; analyzed the models and theory; manuscript writing, ideas, and discussion.

## Footnotes

1 “mountain lion” in Kumeyaay; San Diego State University is built on Kumeyaay land.

2 The original code for this approach is no longer available, and thus `nyemtaay` currently represents the only implementation of this approach.

